# DISEASES 2.0: a weekly updated database of disease–gene associations from text mining and data integration

**DOI:** 10.1101/2021.12.07.471296

**Authors:** Dhouha Grissa, Alexander Junge, Tudor I. Oprea, Lars Juhl Jensen

## Abstract

The scientific knowledge about which genes are involved in which diseases grows rapidly, which makes it difficult to keep up with new publications and genetics datasets. The DISEASES database aims to provide a comprehensive overview by systematically integrating and assigning confidence scores to evidence for disease–gene associations from curated databases, genome-wide association studies (GWAS), and automatic text mining of the biomedical literature. Here, we present a major update to this resource, which greatly increases the number of associations from all these sources. This is especially true for the text-mined associations, which have increased by at least 9-fold at all confidence cutoffs. We show that this dramatic increase is primarily due to adding full-text articles to the text corpus, secondarily due to improvements to both the disease and gene dictionaries used for named entity recognition, and only to a very small extent due to the growth in number of PubMed abstracts. DISEASES now also makes use of a new GWAS database, TIGA, which considerably increased the number of GWAS-derived disease–gene associations. DISEASES itself is also integrated into several other databases and resources, including GeneCards/MalaCards, Pharos/TCRD, and the Cytoscape stringApp. All data in DISEASES is updated on a weekly basis and is available via a web interface at https://diseases.jensenlab.org, from where it can also be downloaded under open licenses.

## 1 Introduction

To understand human diseases at the molecular level, we need a comprehensive overview of which genes are linked to each disease. Since these links can come from many types of data, each of which is growing at a fast rate, there is a need for frequently updated databases that integrate the heterogeneous evidence for disease–gene associations. To this end, we provide the freely available DISEASES database resource [1], which has been continuously updated on a weekly basis since 2014. This resource automatically extracts disease–gene associations from the biomedical literature by identifying mentions of disease and gene names and counting how often they are co-mentioned. These are combined with manually curated associations and experimental evidence from genome-wide association studies (GWAS).

The first version of DISEASES included text mining of 24 million abstracts available from the PubMed database, which has since grown by another 8 million new abstracts. Moreover, text mining of full-text articles can yield approximately 50% more disease–gene associations (at the same false positive rate) than text mining of the corresponding abstracts [2]. Combined with the rapid growth of open-access publishing [3], which allows for text mining and redistribution, this shows a clear opportunity for improving resources like DISEASES to go beyond text mining of abstracts only.

Text mining has been applied to many tasks in the biomedical domain, such as identifying genes and other named entities [4] in text and subsequently extracting associations between genes and other genes [5], pathways [6], and diseases [7]. Many studies have focused on doing the latter based on biomedical abstracts only [8], whereas fewer have included full-text articles too [9]. General text-mining efforts, not specifically focused on disease–gene associations, of course also extract such associations [10, 11].

DISEASES is not the only database to gather evidence of disease–gene associations. Such associations have for many years been manually annotated by curators of both general protein databases, such as UniProtKB/Swiss-Prot [12], and databases focused on genetics of diseases, such as Online Mendelian Inheritance in Man (OMIM) [13] and Medline-Plus from Genetics Home Reference (GHR) [14]. In addition to these broad databases, many specialist databases exist which focus on specific diseases or classes of diseases, including the cancer mutation databases Catalog of Somatic Mutations in Cancer (COSMIC) [15] and intOGen [16].

Genome-wide association studies (GWAS) are another important source of disease–gene associations, which comes with its own ecosystem of database resources. In GWAS, statistically significant associations between single nucleotide polymorphisms (SNPs) and phenotypic traits (including diseases) are identified and used to infer gene–trait associations. These associations, both at the SNP and gene level, are collected by the National Human Genome Research Institute (NHGRI)–European Bioinformatics Institute (EBI) GWAS Catalog [17], GWAS Central [18] databases, and GWASdb [19]. However, GWAS results are complex to interpret; the identified SNPs are typically not the causal mutations, which due to linkage disequilibrium could reside anywhere within a chromosomal region that may contain multiple genes [20]. Several secondary GWAS databases, such as DistiLD [21] and TIGA [22], thus aim to help non-experts interpret GWAS results by integrating other relevant information and prioritizing the results.

Several integrative database resources, like DISEASES, combine many of the types of evidence for disease–gene associations mentioned above. The best known of these are probably MalaCards/GeneCards [23] and DisGeNET [24], which take two very different approaches. The MalaCards and GeneCards databases present one page for each disease and gene, respectively, which gives the user a very comprehensive overview of the available information, including text mining from DISEASES. DisGeNET, on the other hand, has a strong focus on scoring the associations and on making data available amenable to further computational analysis via application programming interfaces (APIs). The latter is also true for knowledge-based aggregators such as Pharos [25] and Open Targets [26], which integrate many types of evidence from numerous sources, including gene–disease associations.

In this paper, we describe the major improvements of the DISEASES resource made since the 2015 publication [1]. The gene set and associated dictionary have been updated to be consistent with the latest version of STRING [27] and the text corpus in DISEASES is now automatically constructed by merging the open-access subset of PubMed Central (PMC) with PubMed abstracts. This has jointly led to substantial improvements of the text-mining results. We have further updated DISEASES to import experimental data from a new GWAS resource, TIGA [22]. We map all disease–gene associations to a common set of identifiers and provide confidence scores for the associations, which are comparable across evidence types. All data are freely available both via a web interface (https://diseases.jensenlab.org/), as bulk download files, and through integration into other resources and tools, specifically Cytoscape, TIN-X, and Pharos.

## 2 Materials & Methods

The DISEASES database combines heterogeneous evidence from several sources. We will go through these, starting with three databases of manually curated disease—gene associations, followed by two sources of experimental evidence, and finally the automatic text mining, which we break down into corpus construction, dictionary construction, named entity recognition (NER), and co-occurrence scoring. Finally, we describe how the DISEASES confidence scores are assigned for each type of evidence.

### 2.1 MedlinePlus

The genetics section of the MedlinePlus resource, formerly known as Genetics Home Reference, includes disease–gene associations obtained from manual curation of the biomedical literature [14]. We first download the list of all diseases and then query the MedlinePlus REST API with each disease to retrieve the list of associated HGNC gene symbols. We then use the dictionaries described later to map the disease names and gene symbols to their Disease Ontology identifiers [28] and STRING v11 identifiers, respectively.

### 2.2 UniProt Knowledgebase (UniProtKB)

The Swiss-Prot section of UniProtKB consists of expert-reviewed protein entries, which include diseases associated with each protein among many other types of biological information [12]. We extract the diseases associated with a protein entry by parsing the keyword field, where they are specified using a controlled vocabulary. We manually mapped these to their corresponding concepts in Disease Ontology and used the dictionary described later to map the UniProtKB accession numbers to STRING v11 identifiers.

### 2.3 Amyloidoses Collection (AmyCo)

The AmyCo database specifically collects information on amyloidoses and other diseases related to amyloid deposition [29]. It contains data from 249 articles on 75 diseases classified into two broad groups: amyloidoses and clinical conditions associated with amyloidosis, including precursors and proteins co-deposited with amyloid deposits. AmyCo identifiers are mapped to their corresponding Disease Ontology identifiers whenever the AmyCo name could be found as an exact synonym; otherwise the AmyCo identifier is mapped to the Disease Ontology broader parent(s).

### 2.4 Target Illumination by GWAS Analytics (TIGA)

TIGA [22] is a new weekly updated web resource that imports GWAS data from the NHGRI-EBI GWAS Catalog [17], maps SNPs to the nearest protein-coding genes, and evaluates the confidence of each gene–trait association. The latter is done by calculating an average rank score based on the number and distance of SNPs supporting the association, the p-value of the most significant SNP, and the number of studies supporting the association weighted by the Relative Citation Ratio [30] of the underlying publications. From TIGA, we extract the subset of traits that are diseases and map their Experimental Factor Ontology (EFO) terms to the corresponding Disease Ontology terms based on ontology cross-references and the EMBL-EBI Ontology Xref Service. The Ensembl gene identifiers are mapped to STRING v11 identifiers using the gene dictionary described later.

### 2.5 Text corpora

As the starting point for doing text mining, a large body of biomedical texts is needed. We compile such a corpus based on the PMC open-access subset [3], which consists of 7.3 million full-text articles, and the PubMed abstract database, which contains 39 million entries, 22 million of which have an English abstract. To construct a combined corpus, we download both PMC and PubMed in XML format, specifically, the BioC version of PMC [31] and the PubMed baseline plus daily updates. The latter updates also mean that retracted articles are automatically removed from the corpus as soon as they have been marked as such in PubMed. As a last step, we exclude 826 publications, which are believed to contain falsified data and to have been created by several recently discovered paper mills [32].

As PMC contains articles in several languages and the text-mining pipeline in DISEASES is designed only for English text, we use the pretrained language detection models from fastText [33] to identify the language of each PMC article and remove articles not in English. We next run the NER software (described later) on the English-language articles to count the number of unique entities found in each article. To eliminate articles that mention long lists of genes or diseases, we removed the 630 PMC articles that mentioned more than 200 genes or diseases. For quality reasons and to have consistent article metadata, we decided to only include articles from PMC that are indexed in PubMed. We thus used the identifier mapping file from PMC to convert PMCIDs to PMIDs, and discarded all PMC articles for which a PMID did not exist. For the remaining articles, we merged the information from PMC and PubMed, using the metadata, title and abstract from PubMed and extending it with the article body text from PMC. Where no PMC open-access version of an article was available, we simply used the metadata, title, and abstract from PubMed.

The corpus in DISEASES is updated every weekend. All results presented in this paper are based on October 20th, 2021 version of the corpus (FullText2021). To assess the impact of including full-text open-access publications, we also have a second corpus (PubMed2021), which includes only the title and abstract text of the same publications. Finally, to be able to assess how much the general growth of PubMed contributes to the performance, we have a third corpus, which consists of ~ 24 million abstracts published by end of 2013 (PubMed2013). This corpus closely resembles what was in the initial version of DISEASES, which was submitted in January 2014 [1].

### 2.6 Dictionaries

For mapping names and identifiers and for recognizing them in text, we need comprehensive dictionaries of human genes and diseases. The dictionary of diseases is constructed based on all the names and synonyms from Disease Ontology [28] and extended with additional amyloidoses from AmyCo, mappings to ICD-10, and manual additions of missing disease synonyms and acronyms. The human gene dictionary was obtained from STRING v11.0 [27] and is based on information from Ensembl [34], UniProtKB [12], and HGNC [35] databases. We further automatically eliminate clashes between HGNC gene symbols and disease names and extend both dictionaries with orthographic variations of names using the exact same rules as in the first version of DISEASES [1].

We frequently update the dictionaries to incorporate changes to Disease Ontology and to correct errors identified by users. In this paper we make use of two frozen versions of the dictionaries, namely one from October 20th, 2021 (Dict2.0) and, for comparison, the dictionaries from the first version of DISEASES [1] (Dict1.0). The latest dictionary is available from the Downloads tab of the DISEASES web resource.

### 2.7 Named entity recognition (NER)

To do NER on the very large text corpora described above — and make frequent updates feasible — a highly efficient tool for matching the dictionaries against the text is needed. As in the previous version of DISEASES, we do this using the Tagger software, which is described in detail elsewhere [36] (https://github.com/larsjuhljensen/tagger). Briefly, the combined dictionary is first loaded into memory in a custom hash table that allows fast, case-insensitive lookup and further allows for arbitrary insertion and deletion of hyphens. We then tokenize the text on white-space and special characters (including hyphen and slash) and look up combinations of tokens in the combined dictionary to identify left-most longest matches. To improve the precision, we globally block tagging of names that would otherwise give rise to many false positives by manually inspecting the tagging results of all names that occur more than 2000 times in PubMed as well as names that gave rise to errors reported by users. Each match in the text is normalized to the unique entity identifier from the dictionary and, in case of diseases, the term is backtracked to all parent terms in Disease Ontology.

### 2.8 Co-occurrence scoring

From the NER results, we calculate co-occurrence score between any given pair of a gene and a disease, which quantifies how much these entities have been mentioned together in the text corpus. The scoring scheme takes into account that co-occurrences within sentences are stronger evidence than cooccurrences across sentences within a paragraph, which in turn are stronger than co-occurrences across paragraphs within a paper. The scoring scheme further takes into account both how much the entities co-occur on an absolute scale and relative to what would be expected by random chance. This approach is that same as was used in DISEASES v1 [1], except that the scoring having been extended to handle full-text articles as previously described [2].

### 2.9 Comparison to DISEASES v1

To facilitate comparison with DISEASES v1, which differs both in terms of the dictionaries and the text corpus used, we generated four datasets of text-mining results. These represent the following combinations of the old and new dictionaries (*Dict1.0* and *Dict2.0*) with text corpora representing the abstracts used in DISEASES v1 (*PubMed2013*), the full set of abstracts now available (*PubMed2021*), and the combined corpus including also full-text articles from PMC (*FullText2021*):

i. *Dict1,PubMed2013*: *PubMed2013* mined using *Dict1.0*, representing the text-mining channel of DISEASES v1 when published
ii. *Dict1,PubMed2021*: *PubMed2021* mined using *Dict1.0* to show the effect of updating DISEASES v1 with new abstracts
iii. *Dict2,PubMed2021*: *PubMed2021* mined using *Dict2.0* to capture the changes attributed to dictionary improvements
iv. *Dict2,FullText2021*: *FullText2021* mined using *Dict2.0*, representing the text-mining channel of DISEASES v2

#### 2.9.1 Gold standard of disease–gene associations

To build an up-to-date gold standard of disease–gene associations, we followed the approach described in the original DISEASES publication [1]. We used the knowledge channel of manually curated annotations imported from UniProtKB and MedlinePlus (doi:10.6084/m9.figshare.17075708). A new reduced benchmarking set is then generated that includes only entity names explicitly annotated in UniProtKB and MedlinePlus and, with diseases broader parent terms. When an annotated disease was a child term of another annotated disease, we kept the broader parent terms and back-tracked the child-term annotations to it via the is a relationships in the ontology. The final gold standard comprises 7,005 of inferred disease–gene associations, all were given a high confidence score of 4 to 5 stars to show how they are biologically meaningful.

#### 2.9.2 Benchmarking of text-mined associations

To evaluate and compare the quality of the text-mining results, we bench-marked each of the four sets of text-mining results (*Dict1.0_PubMed2013*, *Dict1.0_PubMed2021*, *Dict2.0_PubMed2021*, and *Dict2.0_FullText2021*) on the gold standard of disease–gene associations. Given a disease–gene association, we labeled it as positive if the association exists in the gold standard, labeled it as negative if both the disease and the gene (but not the association) exist in the gold standard, and otherwise discarded it. Based on this binary labeling, we constructed receiver operating characteristic (ROC) curves for each of the four sets of text-mining results by sorting the associations descending by score and plotting the true positive rate (TPR) against the false positive rate (FPR). To quantify the difference between the ROC curves, we calculated the area under curve (AUC) for each.

## 3 Results & Discussion

### 3.1 Overview of the DISEASES resource

Figure 1 gives an overview of the disease–gene associations in DISEASES by mapping them onto high-level terms in Disease Ontology that have at least 100 associated genes. For each disease term, the area shows the number of associations with a confidence score of at least 3 stars, which is further broken down based on the source of the associations (knowledge, experiments, or text mining). Automatic text mining is by far the biggest source of associations for all diseases, accounting for more than 60% of the total.

**Figure 1:**
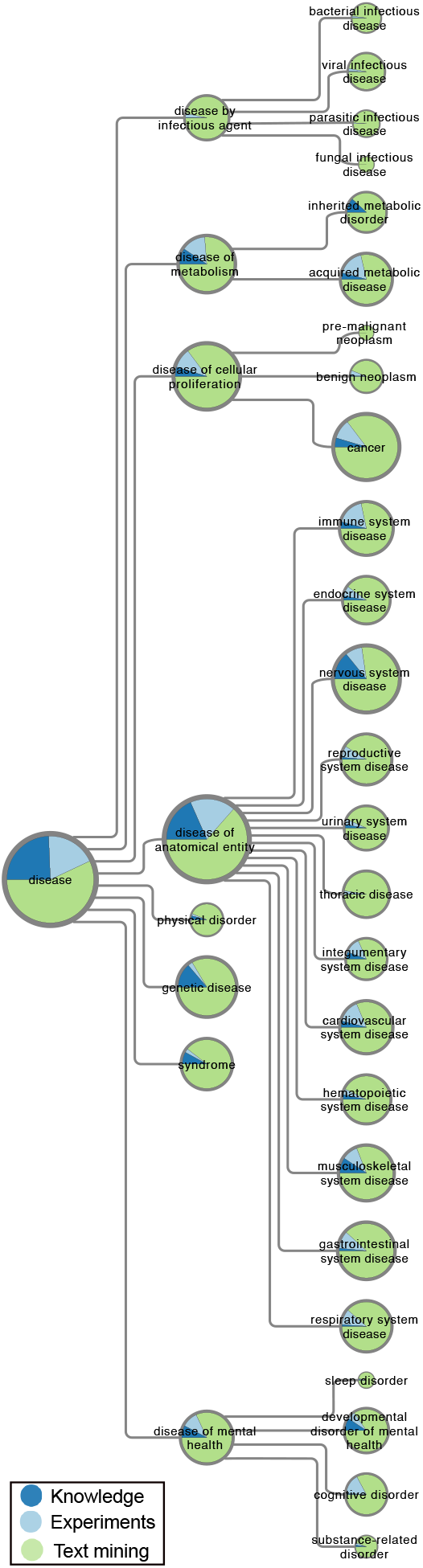
Overview of disease–gene associations in DISEASES. The number of disease–gene associations with a confidence score of at least 3 stars is proportional to the area of the pie charts, which represent high-level terms from Disease Ontology. In each pie chart, the associations are broken down by evidence type, i.e. curated knowledge, GWAS experiments, and automatic text mining of the literature.

The category of diseases with the most associations, especially from knowledge and experiments, is *disease of anatomical entity*, within which we see a fairly even distribution across many anatomical systems. This is followed by *disease of cellular proliferation*, which almost exclusively covers cancer–gene associations. By contrast, we find quite few associations (7443) for *disease of infectious agent*, which come exclusively from text mining.

### 3.2 Growth of the number of disease-gene associations

The number of open access articles available from PMC has been grown exponentially over time, reaching more than 7 million as of September 2021. The inclusion of these — as well as the more than 8 million new PubMed abstracts published since the initial release of DISEASES in 2015 — has an obvious and direct effect on the number of disease–gene association one can find by text mining. While the text-mining channel in the DISEASES database is our main focus in this article, it is not the only improvement. Table 1 provides an overview of the content of the original and the new releases DISEASES database, showing the number of genes, diseases, and associations provided by each evidence channel.

**Table 1:** Comparison of the new and original versions of DISEASES. For each evidence channel we show the number of associations, unique genes, and unique diseases for both the new and the originally published version of DISEASES. In case of the *knowledge* and *experiments* channels, these numbers are further provided for each of the source databases. The numbers in parentheses are the counts before evidence was backtracked to parent terms in Disease Ontology. For the *text mining* channel, we instead subdivide the counts by confidence score.

| Evidence channel | Associations |  | Genes | Diseases |  |
| --- | --- | --- | --- | --- | --- |
| Knowledge |  |  |  |  |  |
| DISEASES 2021 | 36,448 | (6,739) | 3,723 | 1,528 | (1,048) |
| AmyCo | 1,225 | (186) | 57 | 180 | (52) |
| MedlinePlus | 23,957 | (3,516) | 2,383 | 1,460 | (966) |
| UniProtKB | 19,517 | (3,739) | 2,642 | 270 | (119) |
| DISEASES 2015 | 15,231 | (2,953) | 2,001 | 735 | (453) |
| GHR | 7,551 | (1,169) | 965 | 671 | (390) |
| UniProtKB | 11,576 | (2,187) | 1,651 | 271 | (120) |
| Experiments |  |  |  |  |  |
| DISEASES 2021 | 152,611 | (26,346) | 9,180 | 574 | (295) |
| TIGA | 152,611 | (26,346) | 9,180 | 574 | (295) |
| DISEASES 2015 | 89,073 | (20,206) | 10,711 | 423 | (264) |
| COSMIC | 55,791 | (13,050) | 8,786 | 142 | (76) |
| DistiLD | 36,650 | (7,185) | 4,315 | 351 | (210) |
| Text mining |  |  |  |  |  |
| DISEASES 2021 | 4,512,870 |  | 19,116 | 8,537 |  |
| 4 star confidence | 18,129 |  | 2,988 | 2,959 |  |
| 3 star confidence | 224,642 |  | 11,207 | 6,711 |  |
| 2 star confidence | 1,659,331 |  | 18,913 | 8,342 |  |
| DISEASES 2015 | 478,407 |  | 15,631 | 4,598 |  |
| 4 star confidence | 1,044 |  | 478 | 662 |  |
| 3 star confidence | 15,226 |  | 3,207 | 2,267 |  |
| 2 star confidence | 142,892 |  | 12,706 | 4,354 |  |

The *Knowledge* channel has more than doubled in terms of both disease– gene associations and unique diseases covered. This growth comes primarily from GHR, which has in the meantime been integrated into MedlinePlus, but UniProtKB has also grown substantially. AmyCo contributes a comparably low number of new associations, since it covers only a specific type of diseases.

The *Experiments* channel has in many ways between the two versions. Replacing DistiLD with TIGA has increased the number GWAS-based associations by more than a factor of four and more than doubled the coverage of genes. However, with the new release we have also to remove COSMIC for license reasons, thus losing more than half of the experimental associations in the original release of DISEASES. All in all, the *Experiments* channels has grown by over 70%.

For the *Text mining* channel, we subdivide the evidence by their confidence scores as presented in the DISEASES web interface. That is, we present the number of genes, diseases, and associations that rated as 2-, 3-, and 4-star confidence as well as the total numbers for the channel (including association scoring below 2 stars). The number of associations has increased dramatically at all confidence levels and especially at high confidence levels, with over 9-fold increase overall and over 17-fold increase for 4-star associations. The same trend holds true when looking at the numbers of unique genes and diseases covered.

### 3.3 Improved quality of text-mined associations

We assessed the quality of the disease–gene associations from the new version of DISEASES to the originally published version by benchmarking both against a gold standard of manually annotated gene–disease associations (see Materials & Methods for details). The results are shown as ROC curves in Figure 2, which reveals a substantial improvement both overall (AUC increasing from 0.829 to 0.916) and in the low false-positive-rate part, which is arguably the most relevant part. As the ROC curve for the new version is consistently above that of the original one, the new version constitutes an improvement regardless of whether the user cares most about getting higher true positive rate or lower false positive rate.

**Figure 2:**
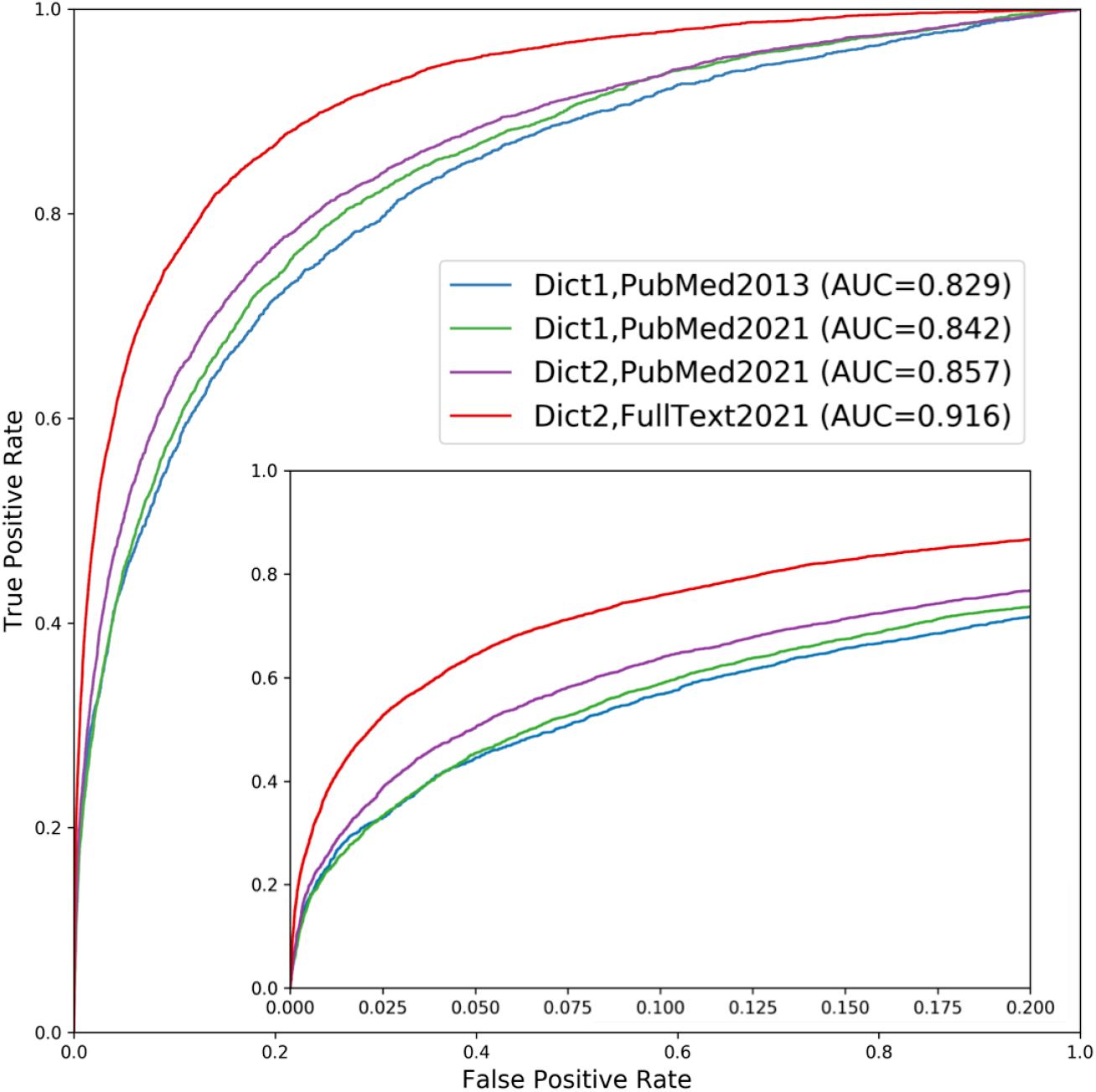
Performance improvement of the *text mining* channel. As shown in the receiver operating characteristic (ROC) curves, text mining performs markedly better in the new version of DISEASES (Dict2,FullText2021) compared to the originally published one (Dict1,PubMed2013). To quantify the sources of improvements, we show two additional curves: one using the new dictionary on the latest abstract collection only (Dict2,PubMed2021), and another using the old dictionary on the same abstracts (Dict1,PubMed2021). Comparing the curves reveals that most of the improvement stems from the addition of full-text articles, but that the new disease and gene dictionaries also led to considerable improvement. By contrast, the growth in PubMed abstracts from 2013 to 2021 made only a minor difference. The insert shows a zoom of the high-confidence part of the plot.

This performance improvement is due to a combination of i) general growth in the number of biomedical abstracts available from PubMed, ii) improvements to the dictionaries used for NER, and iii) the addition of full-text articles from the PMC open access subset. To quantify the importance of each of these factors, we show two additional ROC curves in Figure 2: performance when updating with new abstracts but still using the original dictionaries and performance when further updating the dictionaries. Comparing the four ROC curves shows that the growth of PubMed abstracts alone gives only a small improvement of the AUROC from 0.845 to 0.859. The use of the new dictionaries leads to a bigger incremental improvement, increasing the AUROC from 0.859 to 0.866. However, the addition of full-text articles to the corpus is responsible for the biggest improvement, bringing the AUROC from 0.866 to = 0.922.

These results show that while the growth of the literature does give an almost free improvement, only requiring the pipeline to rerun on latest PubMed, the vast majority of the improvement seen between the original version of DISEASES and the new version stems from our work on improving the dictionaries used for NER and on integrating full-text articles into the corpus. The results also highlight how important it is for text mining efforts to be permitted to process full-text articles rather than only abstracts.

### 3.4 Research paper mills

To the best of our knowledge, all assessment of text-mining results to date have focused purely on the ability of a text-mining system to correctly extract what is stated in the text. However, from the perspective of using text mining to construct a knowledgebase from literature, is equally important if what is stated in the text is true. Co-mentioning-based systems indirectly address this, since high-scoring associations will be supported by multiple publications.

Recently, the problem of incorrect information in the literature has become a bigger concern due to the discovery of so-called “paper mills”. These appear to be companies that mass produce fake articles and sell them to researchers at Chinese hospitals [37]. As these articles were published in international journals indexed in PubMed, they would by default be included in our text corpus, thus providing false support for disease–gene associations in our database. To avoid this, we have compiled a list of the 826 papers identified so far to originate from paper mills (as per June 2021) and explicitly exclude these from our text corpus. To allow others to also easily exclude these papers, the latest list is available for download from the DISEASES website.

### 3.5 Integration into other resources

Just like DISEASES itself builds upon other databases, we have designed it to be easy to integrate into other resources. We do this both from a technical perspective by providing simple bulk download options and from a legal perspective by not integrating any data that would prevent us using an open license. Several tools and databases already take advantage of this, importing either disease–gene associations from all evidence channels or specifically the associations from text mining.

The GeneCards and MalaCards databases, both members of the GeneCards suite, provide a comprehensive overview of information on human genes, including diseases associations, by integrating evidence from 150 sources [38]. One of these is the text-mined disease–gene associations from DISEASES, which GeneCards/MalaCards downloads on a regular basis and combines with associations from other source databases. The GeneCards/MalaCards web resources link back to DISEASES website to allow users to easily inspect the text-mining evidence for any given association. The Target Central Resource Database (TCRD) and the associated Pharos [25] and TIN-X web resources [39], which aim to shed light on potential new drug targets, similarly obtain up-to-date disease–gene associations from the DISEASES database. DISEASES (and “tagger” output) are also an integral part of Geneshot [40] and Harmonizome [41].

DISEASES is also designed to interface easily with the STRING, COMPARTMENTS and TISSUES resources by using the same gene identifiers. Through the Cytoscape app *stringApp*, it is thus possible to quickly retrieve a human protein network for any disease of interest [42]. To do this, stringApp first queries the DISEASES database to obtain a list of genes associated with the disease and subsequently queries STRING to obtain the corresponding protein network.

## 4 Conclusion

The DISEASES database has since 2014 provided the community with a weekly updated resource of disease–gene associations. The latest version features several important improvements compared to the original publication. In addition to text mining PubMed, which has meantime grown by another 8 million abstracts, the DISEASES text corpus now also includes open access full-text articles from PubMed Central. Together with technical improvements to the text-mining pipeline itself, this has led to a > 9-fold increase in the number of disease–gene associations extracted at any confidence cutoff. DISEASES has also been upgraded to use GWAS data via the new TIGA database [22], which increased the number of experimental associations by more than 70%.

The database is freely available at https://diseases.jensenlab.org/ where it can be browsed via a web interface as well as downloaded in its entirety to facilitate large-scale analysis. Moreover, DISEASES is designed to integrate easily with other resources, and the disease–gene associations are available through other resources, including the GeneCards/MalaCards and TCRD/Pharos databases, Harmonizome, Geneshot, and the Cytoscape stringApp.

## Acknowledgements

This work was supported by US National Institutes of Health grant U24 224370 for “Illuminating the Druggable Genome Knowledge Management Center” (IDG KMC), and by the Novo Nordisk Foundation (grant number NNF14CC0001).

